# Automated design of gene circuits with optimal mushroom-bifurcation behaviour

**DOI:** 10.1101/2022.05.09.490426

**Authors:** Irene Otero-Muras, Ruben Perez-Carrasco, Julio R. Banga, Chris P. Barnes

## Abstract

Recent advances in synthetic biology are enabling exciting technologies, including the next generation of biosensors, the rational design of cell memory, modulated synthetic cell differentiation and generic multi-functional bio-circuits. These novel applications require the design of gene circuits leading to sophisticated behaviours and functionalities. At the same time, designs need to be kept minimal to avoid compromising cell viability. Bifurcation theory of dynamical systems provides powerful tools to address complex nonlinear dynamics and multifunctionality, linking model topology and kinetic parameters with circuit behaviour. However, the challenge of incorporating bifurcation analysis to automated design has not been accomplished so far. In this work we present an optimisation-based method for the automated forward design of synthetic gene circuits with specified bifurcation diagrams, allowing us to find minimal topologies optimizing the required functionalities and taking into account additional requirements and/or context specifications. We apply the method to design of gene circuits exhibiting the so called mushroom bifurcation, a relatively unexplored multi-functional behaviour of particular relevance for developmental biology. Using the results of the optimisation analysis we explore the capabilities of the resulting circuits for possible applications in advanced biosensors, memory devices, and synthetic cell differentiation.

## 1 Introduction

Real-world and cutting edge applications of synthetic biology are demanding circuit designs with increasingly complex behaviours. Towards building the synthetic cell from the bottom-up, *new developments are expected*, quoting Schwille et al. (2018), *when designing cellular systems featuring complex behaviors, including division, cognition, and motility*.

One of the main challenges of synthetic biology is to design and implement gene regulatory circuits capable of complex behaviours in a near-optimal fashion while keeping a minimal design (Schaerli et al. 2014, Pérez-Carrasco et al. 2018). A milestone in gene circuit automated design is CELLO (Alec A. K. Nielsen et al. 2016) enabling the design of circuits with pre-specified steady state and input-output behaviours, and for the first time proving good predictability in living cells of model-based automated design software. Aiming to address more complex (and dynamic) behaviours, tools based on Mixed Integer Nonlinear Programming have shown good flexibility and computational efficiency (Otero-Muras et al. 2016, Otero-Muras & Banga 2017).

The limited resources of the cell restrict the combination of multiple working circuits in the same organism. This gives leading relevance to the design of multifunctionality, regulatory networks capable of distinct dynamical behaviours. But, how can different behaviours be integrated in the same circuit? How to endow a cell with the capacity to respond differently to a signal leading to complex dynamical behaviours? Bifurcation theory of dynamical systems provides powerful tools to answer these questions, enabling the mapping between the topology of the network (given by a set of parametrized differential equations) and the different dynamics available under a controllable input or signal. Each bifurcation of the system changes the number and/or nature of the long-term dynamics of the system, e.g. from monostable to bistable, or from a stable steady state to an oscillator. These phenomena can be represented by bifurcation diagrams, showing how the number, position, and dynamics of each steady state change under a set of controllable parameters. However, standard tools for bifurcation analysis are based on continuation algorithms which require precise a priori knowledge on parameters and steady state solutions, hampering the integration of bifurcation diagrams within automated algorithms for circuit design. Alternative methodologies based on chemical reaction network theory (Yordanov et al. 2020, Reyes et al. 2020, Otero-Muras & Banga 2021) have paved the way for the integration of bifurcation theory into the automated design of biocircuits, an ambitious design framework which has not been addressed so far.

In this work we address this last topic and present a method for automated design of gene circuits with pre-specified bifurcation diagrams. The method extends and combines efficient global mixed-integer nonlinear optimization methods with an innovative procedure for bifurcation detection, allowing the following novel features:

i. Automated design of gene circuits that not only exhibit behaviours compatible with a target bifurcation diagram, but are also optimized for specific additional criteria (like metabolic cost) given by sets of functions.
ii. Systematic exploration of minimal topologies compatible with the required behaviour.
iii. Robust design i.e. finding robust topologies with respect to parameter perturbations (i.e. topologies that remain functional in spite of perturbations in the parameters).
iv. More sophisticated design tasks, computing optimal trade-offs between design objectives and finding Pareto optimal topologies with respect to different opposing criteria (such as performance and metabolic cost, or robustness and topological complexity).

A multifunctional behaviour of special interest for synthetic biology applications is the mushroom bifurcation, named after its mushroom-shaped bifurcation diagram, originally identified in the differentiation of neural stem cells (Song et al. 2006, Sengupta & Kar 2018, Giri & Kar 2021). The mushroom can result from the combination of two toggle switches, giving rise to four saddle node bifurcations (Fig. 1). Similarly to the toggle switch, the mushroom presents two steady-states, but in contrast, one of the steady states (termed ON state) is only available for a window of intermediate values of the signal, while the other (termed OFF state) is available for high and low values of the signal. Therefore, in certain aspects, the phenotypic behaviour of the mushroom is an extension of ‘band-detect’ gene regulatory networks, which are constructed from incoherent feed forward loops (Basu et al. 2005, Sohka et al. 2009). These have recently been applied to model French flag type pattern formation (Schaerli et al. 2014) and other programmed spatial behaviour (Kong et al. 2017). The mushroom-shaped locus of equilibria, leading to two different ranges of bistability provides the system with unique hysteresis properties where the state of the cell will be determined by the signal history. Bistability (and multistability in general) is in itself a property of interest in synthetic biology, microorganisms, and mammalian cells (Gardner et al. 2000, Litcofsky et al. 2012, Zhu et al. 2021), forming the basis of memory, cell decision making systems, biological computation, and pattern formation (Verd et al. 2014, Dalchau et al. 2018, Perez-Carrasco et al. 2016, Barbier et al. 2020). In addition, we can explore the phenotypic space close to the mushroom bifurcation. In particular mushroom topologies can also lead to isola bifurcation diagrams, which, though well-known in dynamical systems theory, has never been demonstrated experimentally. Together, these properties demonstrate how combining common motifs gives rise to an emergent range of dynamical behaviours not described by the individual components, a growing area of interest in systems biology (Jiménez et al. 2017, Pérez-Carrasco et al. 2018). Hence the contribution of this study is twofold: on the one hand incorporating for the first time target bifurcation behaviours in the automated design of biocircuits, while on the other exploring the capabilities of the identified robust mushroom topologies showing how automated exploration can be connected to functional design.

**Figure 1:**
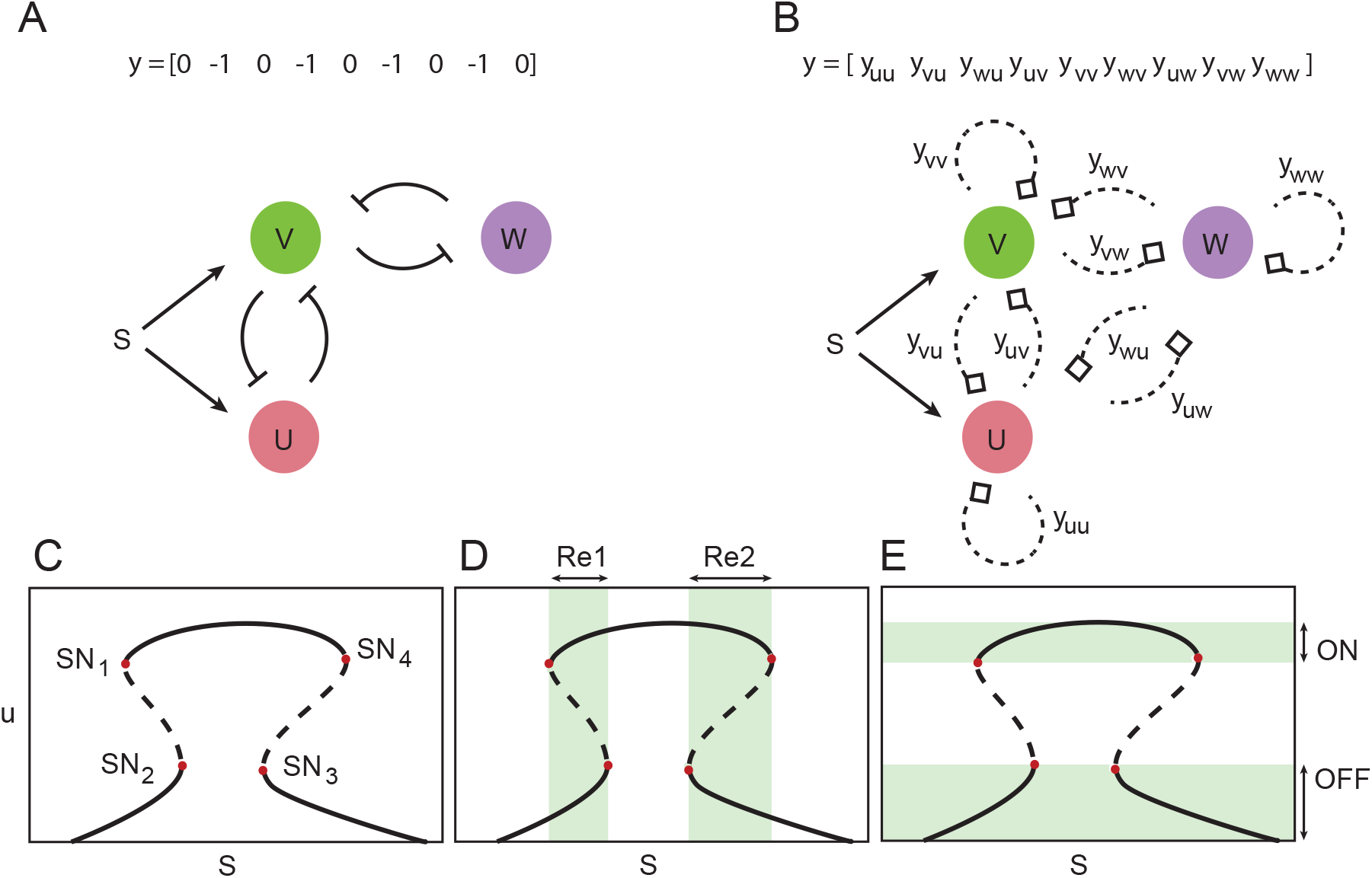
Searching for gene circuit topologies leading to mushroom bifurcation behaviour. A) Gene network containing two pairs of cross-repressing genes. B) The circuit super structure employed in this work to search for mushroom behaviour: super graph and vector encoding. C) The mushroom bifurcation diagram shows four saddle-node bifurcations (indicated by red dots), D) two different bistability ranges (Re1, Re2) and E) ON and OFF levels (shadowed).

## 2 Results

### 2.1 Minimal topologies leading to mushroom bifurcation

Given that the mushroom bifurcation diagram contains two saddle-node bifurcations, we expect that a network containing two pairs of cross-repressing genes (Fig 1 A) can give rise to the target bifurcation diagram. Nevertheless we hypothesized that simpler topologies with less connections can also yield the same functionality. In order to explore this idea we defined the superstructure in Fig 1 B. Starting from this superstructure, which includes 3 genes (U, V, W) with a signal S inducing genes U and V, we looked for circuits with the mushroom characteristic bifurcation diagram. There are 9 potential connection arrows between the nodes (genes) represented by a vector *y* of integers, such that *y*_*ij*_ = − 1 if gene *j* inhibits gene *i, y*_*ij*_ = +1 if gene *j* activates gene *i* and zero otherwise. The mushroom characteristic bifurcation diagram is depicted in Fig 1 C, with 4 saddle-node points (SN) delimiting the steady-state branches, and generating two bistability regions (Re1 and Re2 in Fig. 1 D). This results in an ON state available only for intermediate values of the signal (see Fig. 1 E). We search for topologies leading to a mushroom bifurcation diagram behaviour using an optimization algorithm and a multistart strategy (as described in the Methods section). In order to explore minimal topologies leading to the target behaviour, we first impose a number of 2 genes and a number of 2, 3 and 4 connections.

From the potential 72 connected topologies, only 7 topologies are found leading to mushroom behaviour, represented in Fig. 2 and classified attending to the number of active connections. From this set of topologies, there are three core topologies (for which no connection can be removed without losing the mushroom functionality), corresponding to structures A2, A5 and A7. Surprisingly, the minimal structure leading to a mushroom bifurcation (A2) only requires two nodes, resembling the paradigmatic cross-repressing topology encoding the bistable switch (Gardner et al. 2000). The key distinguishing feature with this topology is the requirement of the signal activating both genes. A screening looking for the existence of mushroom bifurcation diagrams in the presence of a single activating input did not return any successful results, suggesting that the double activating role of the input is a requirement to reproduce the mushroom diagram.

**Figure 2:**
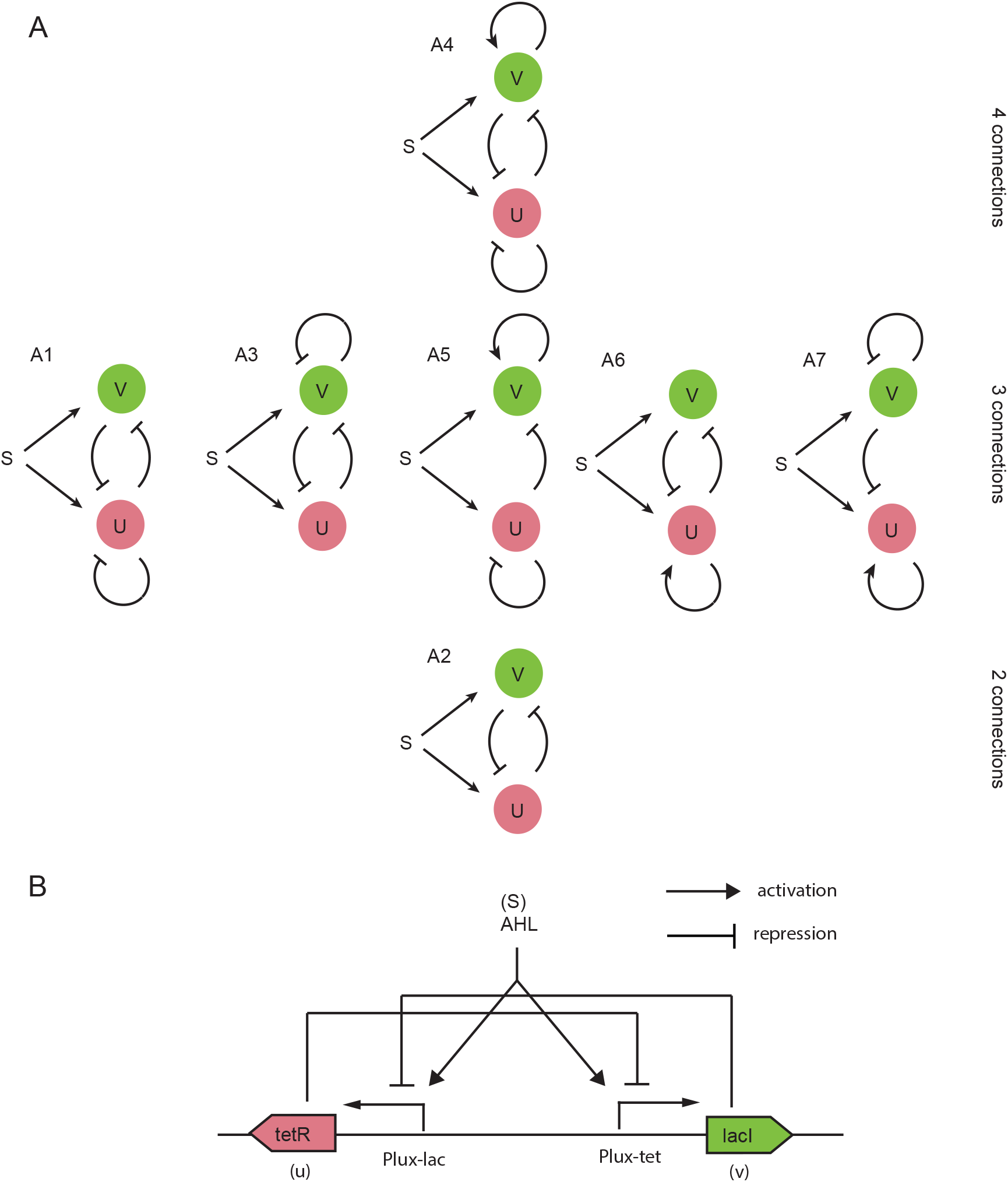
Gene regulatory circuits with the capacity for mushroom bifurcation behaviour. A) All the 2-gene structures found by the algorithm leading to mushroom bifurcations, labelled according to their frequency within the set of solutions (A1, A7 are the most and least frequent, respectively). The topologies are labelled according to their frequency within the set of solutions (*Mushroom2D*). Parameter bounds for the search are included in supplementary Table S1. B) One possible synthetic realization of the circuit with minimal topology (A2) leading to mushroom bifurcation.

With the same optimization strategy, we explore three-dimensional topologies leading to mushroom bifurcations, by fixing in this case the number of genes to 3. The corresponding bounds for the parameters for the 3-gene network are included in the supplemental Table S2. The algorithm detected through multiple optimization runs, more than 300 different 3 gene topologies leading to mushroom bifurcations from the potential 1728 topologies (the set of solutions we denote as *Mushroom3D*). Unlike exhaustive exploration strategies, our optimization-based approach finds structures fulfilling the target behaviour very efficiently, in the order of seconds per run using a standard PC. The most frequent structures found are depicted in the supplemental Table S3 and Figure S1. A selection of 3-gene structures which are not built up from 2-gene mushroom topologies are illustrated in the supplemental Tables S4, S5 and the supplemental Figure S2.

### 2.2 Robust functionality vs topological complexity

Robust design is a recurrent topic in engineering (Mayne et al. 1982), and it has particular relevance in synthetic biology. Usually synthetic biology applications demand designs that are functional (i.e. show the target function or behaviour) in scenarios of high parametric uncertainty, due to potential imprecise characterization of parts and systems, context related parameter variations, etc. (Lormeau et al. 2016). Handling uncertainty is a challenge, and important efforts are taken in this direction (Woods et al. 2016, Asmus et al. 2017, Otero-Muras & Banga 2019, Rybiński et al. 2020).

Here we denote the circuit as functional if it shows a mushroom bifurcation, and we assess its robustness by quantifying how the circuit functionality is kept with respect to perturbations in the parameters.

We define a proxy to assess the robustness of the structures, extending the ideas outlined by Otero-Muras & Banga (2019). The proxy is based on the interquartile ranges (IQR), and measures how large is the spread of the parameter distribution under which the topology is functional as a mushroom. The most robust structures are shown in Fig. 3 A. The structures found (B1 to B384) are labelled according to their frequency within the solution set. From the point of view of circuit implementation, we are interested in finding a good compromise between robustness and the number of connections. In order to select the best structures we build a Pareto front in the objective space (robustness vs number of connections) as depicted in Fig. 3 B. The structures in the Pareto front are shown in Fig. 3 C.

**Figure 3:**
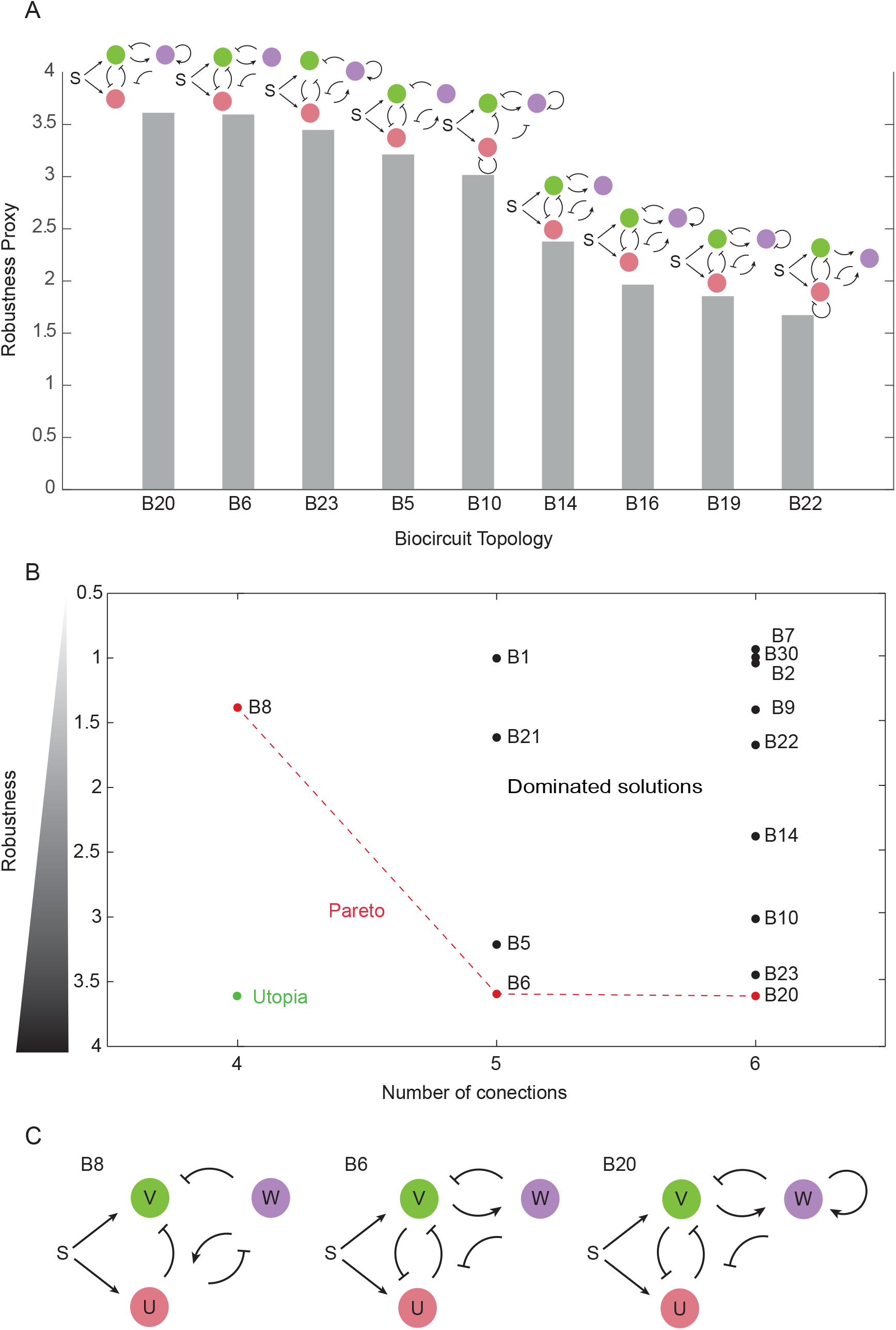
A robustness analysis is performed to find the topologies for which the mushroom functionality is more robust with respect to perturbations in the parameters. A) Proxies for the most robust 3-gene topologies. B) Pareto front of topologies that optimally trade-off robustness and complexity (in terms of the number of connections). C) Structures B8, B6 and B20 optimally trade-off robustness and complexity.

### 2.3 Mushroom annihilation and isola formation

Following the analysis of the mushroom robustness, we explored the resulting bifurcation diagrams when the mushroom is lost. We observed three main transitions out of the mushroom bifurcation diagram (*M*_*1*_ in Fig. 4 A): First, the mushroom head can cross values of saturation or absence or signal, resulting in incomplete mushrooms (*M*_*2*_,*M*_*3*_). Second, a pair of saddle-nodes on one side of the mushroom collide, giving place to a cusp bifurcation giving raise to a bifurcation diagram similar to a bistable switch (*B*). Finally, the two saddle-nodes forming the neck of the mushroom can collide, pinching the neck of the mushroom and producing an isola (*I*_*1*_), a closed curve of equilibrium solutions delimited by two saddle nodes (Fig. 4 B). Strikingly, all these transitions are observed without the requirement to change the topology of the system, making this rich dynamical scenario exploration attainable in gene regulatory circuits.

**Figure 4:**
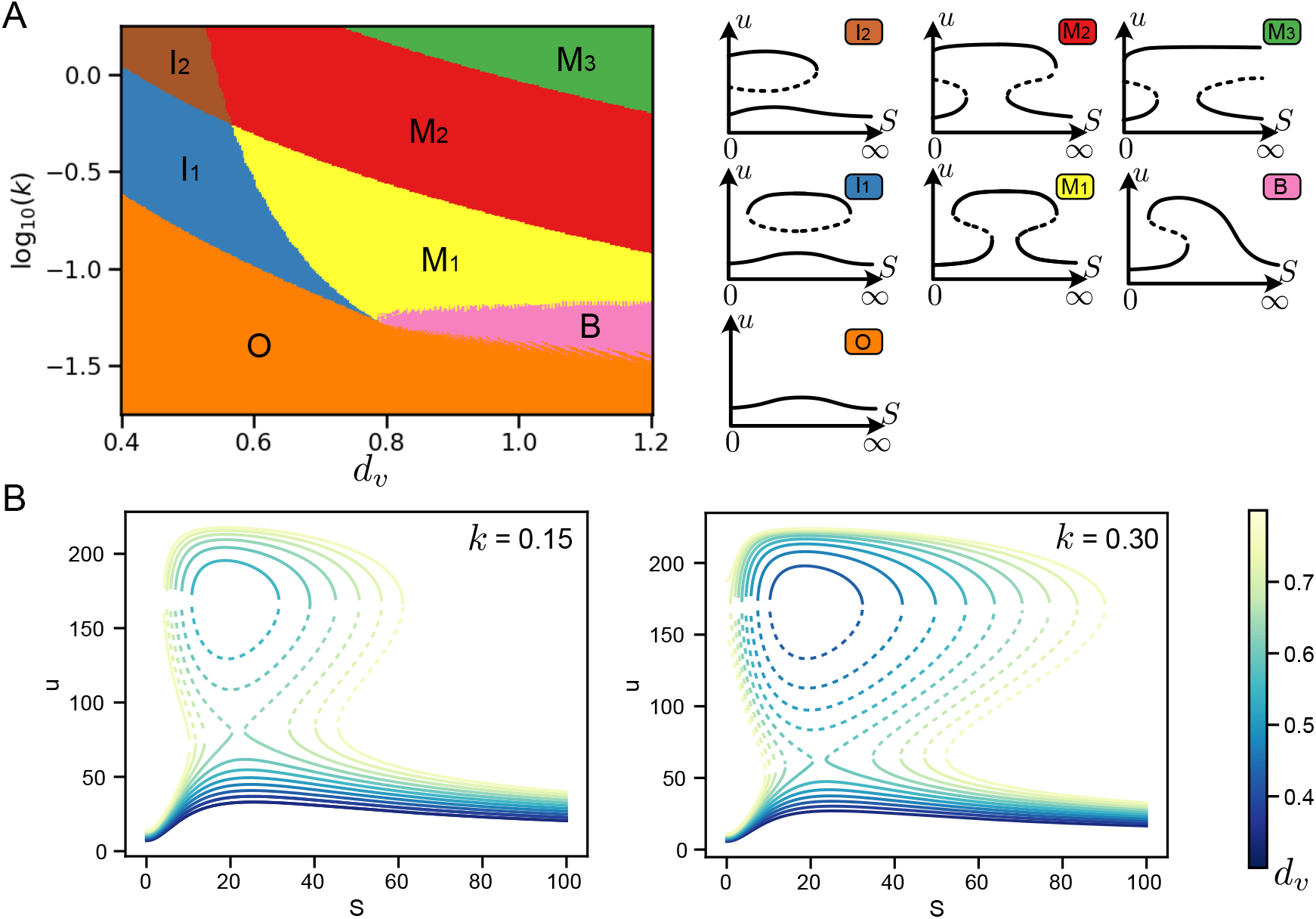
Bifurcation diagrams surrounding the mushroom behaviour reveal a controllable rich functional landscape. A) Phase diagram around a mushroom generated by varying the degradation rate *d*_*v*_ and the interaction strength *k*. Different colors show different bifurcation diagrams. A schematic of each bifurcation diagram is shown in the right panel indicating the different continuation combinations of stable (solid lines) and unstable states (dashed lines). B) Change in the mushroom bifurcation diagram by changing the degradation rate *d*_*v*_ for two different values of *k*. The different diagrams show the annihilation of the mushroom into an isola, followed by the collapse of the isola leading to monostability. Parameters used are: *p*_1_ = 50, *p*_2_ = 1000, *R*_1_ = 264, *R*_2_ = 275, *K*_1_ = 10, *K*_2_ = 133, *d*_*u*_ = 1 for topology A3.

While all the landscapes provide different dynamical properties, isolas are particularly compelling for their cell fate decision capabilities. Isolas are found in different contexts including chemical reactors, buckling of elastic shells and plasma physics (Dellwo et al. 1982, Avitabile et al. 2012). Mechanisms of birth and anihilation of isolas have been classified via singularity theory (Avitabile et al. 2012), being one of such mechanisms with particular interest the transformation a of a mushroom to an isola by reducing the number of saddle-node bifurcations Ganapathisubramanian & Showalter 1984), Song et al. 2006). It is interesting that despite being found both theoretically and experimentally in chemical systems, the isola bifurcation diagram has not yet been directly observed in a biological system. Understanding how to construct such a system in a gene network could therefore enable experimental verification of the underlying dynamical theory. In addition, as we will see below, the isola can form the basis of permanent switching systems, allowing new functionalities not achievable with a genetic toggle switch.

Building a genetic regulatory network with a functional isola requires to be able to prescribe the range of signals of the isola. In our parameter exploration we observed a controllable diversity on the level and range of signals delimiting the isola. In addition, we also detected different degrees of robustness as parameters are varied (Fig. 5B). In order to understand how prevalent is the formation of the isola we explored the formation of isolas in all the identified mushrooms 2D topologies and parameter sets by varying the degradation rate of the node V (*d*_*v*_) and keeping the rest of parameters constant. Strikingly, all the topologies tested exhibited the formation of isolas suggesting that the isola formation is a robust property of the mushroom bifurcation.

**Figure 5:**
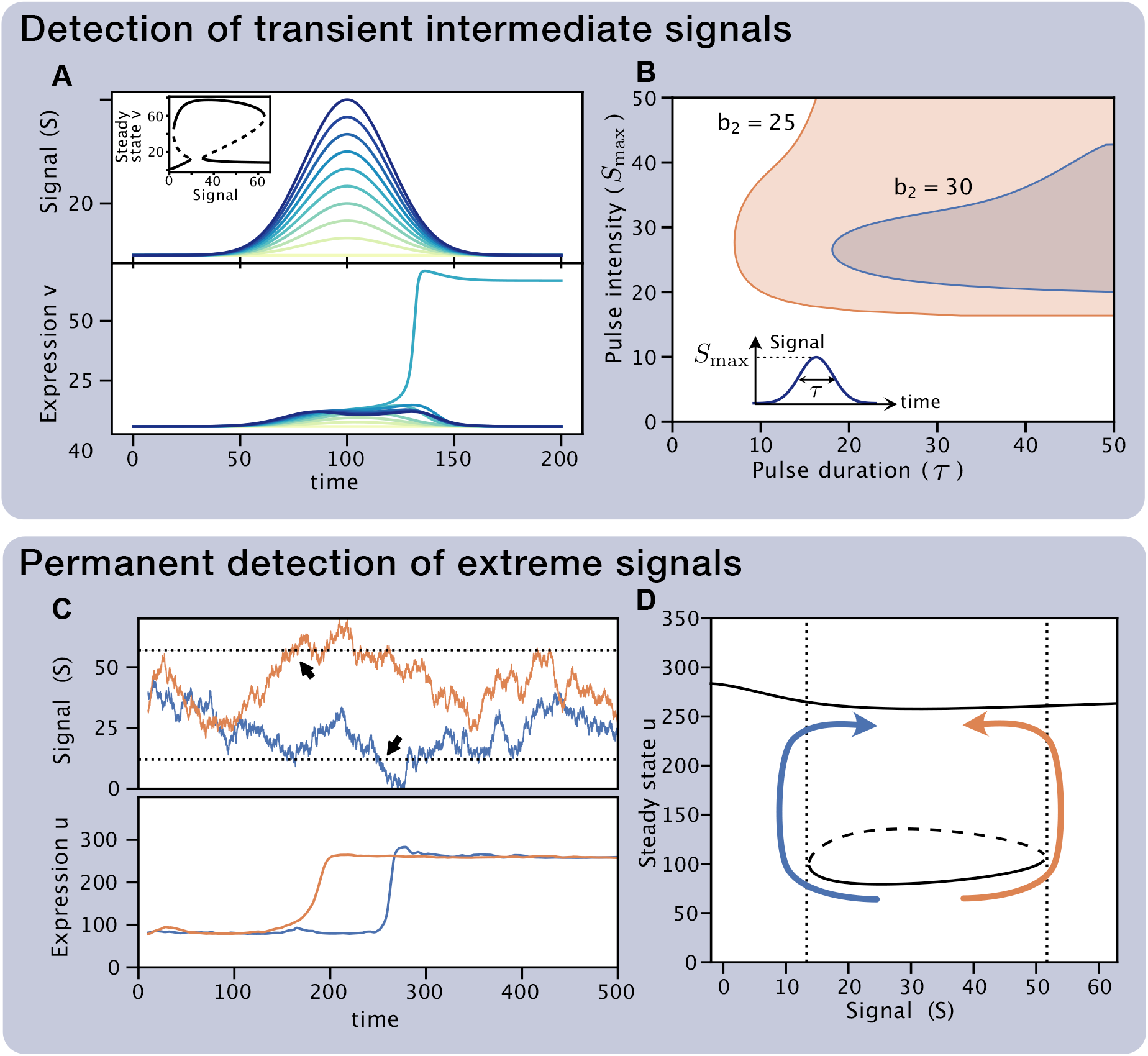
The mushroom circuit is able to discern intensities and durations of transient signals (top). It can also be used to design a sensor of extreme signal values with memory using the isola regime (bottom). A) Response of the mushroom (inset) for different temporal signal pulses (top) of the same duration (*τ* = 20) but different intensity (indicated by color). Expression of gene v (bottom) only becomes stably activated for pulses of intermediate maximum intensity (*S*_max_). B) Response of the mushroom to signal pulses of different duration *(τ)* and maximum intensity (*S* _max_) for two different values of the parameter *b*_2_. Shaded regions indicate combinations of parameters for which node v is activated stably. Signal profiles follow the shape 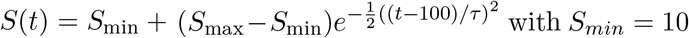. Results correspond to topology A3 with parameters *p*_0_ = 297, *p*_1_ = 30, *p*_2_ = 261,*k* = 9.04 · 10^2^, *R*_1_ = 299, *R*_2_ = 143, *d*_*u*_ = 1, *d*_*v*_ = 0.722, *K*_1_ = 137, *K*_2_ = 10.0. C) Expression of the circuit (bottom) for two different realizations of a noisy signal (top). D) Isola bifurcation diagram showing the detection mechanism. When the signal reaches high or low levels determined by the isola boundary (dotted lines and black arrows in C and D), the system changes steady-state irreversibly. Schematic of the irreversible transitions is indicated by colored arrows. Results correspond to topology A3 with parameters *p*_0_ = 297, *p*_1_ = 200, *p*_2_ = 261,*k* = 9.04 · 10^2^, *R*_1_ = 299, *R*_2_ = 143, *d*_*u*_ = 1, d_*v*_ = 0.722, *K*_1_ = 137, *K*_2_ = 10.0. Initial state was set by opening the mushroom setting parameter *p*_1_ = 20. Noisy signal is a Wiener process.

### 2.4 Design of biosensors and memory devices

The bifurcation diagrams identified in the manuscript are unexplored in synthetic biology and open the door to new functionalities. One of the main characteristics of the mushroom is that the ON state is only available for a reduced range of signals (the neck of the mushroom). This reveals the mushroom capabilities as a biosensor, only allowing the activation (transition from OFF to ON) for precise levels of a target signal located in the neck of the mushroom. Furthermore since the neck of the mushroom is surrounded by two saddle-node bifurcations, the resulting dynamical ghosts will slow down the transition to the ON state (Perez-Carrasco et al. 2016) Hence, activation is not immediate, filtering out transient signals, requiring the persistence of the target signal for a certain amount of time in order to reach the ON state (Fig. 5) A). In addition, the size of the head of the mushroom provides memory of the activation, preserving the an ON state for a larger range of signals than the ones required for the activation (Fig. 5 A). As discussed in the previous section the mushroom bifurcation diagram is prone to pinching by controlling parameters such as the degradation rate of one of the genes. This provides a mechanism to regulate the size of the neck mushroom, opening the door to controlling the range of target signals as well as a the required duration of such signals. This way the mushroom not only serves an accurate signal detector but also as a timer (Fig. 5 B).

In addition to the mushroom bifurcation, the isola also provides compelling functionalities. In contrast to a genetic toggle switch, transitions out from the isola are irreversible. Hence, once in the isola state the system is able to detect a signal that goes outside of the range of the isola (either low or high), unable to reach back the isola independently of future levels of the signal (Fig. 5 C and D). Thus the isola can serve as a sensor of extreme values with infinite memory, detecting if the signal has been high or low at any time point in the past. Since the isola bifurcation diagram is close to the mushroom diagram, reset of the system onto the isola state can be done by “opening” and “closing” the neck of the mushroom.

## 3 Discussion

In this work we have developed an approach to biosystem design based on the specification of bifurcation structure and applied it to the case of the mushroom bifurcation of interest in developmental and synthetic biology. We found gene regulatory networks composed of two and three genes that displayed the desired behaviour. In particular we found that a system based on the genetic toggle switch incorporating a signal activating both genes was sufficient to reproduce a mushroom bifurcation diagram. We also explored the robustness of the these circuits and built a Pareto front to capture the trade-off between the robustness and number of connections, identifying the best topologies to implement the circuit using synthetic biology tools. In addition, we explore specific dynamical functionalities of the mushroom that can be exploited to build biosensors with tuneable temporal and precision properties. We extended our exploration to the formation of different bifurcation diagrams, which occur in the parameter space close to the mushroom bifurcation. We showed that a variety of dynamical behaviours can be created that can provide interesting memory and signal detecting capabilities unattainable with previous memory devices. Interestingly this variety of behaviours was attained mainly by inducing differences in the degradation rates of the genes of the network, which is a common perturbation tool in synthetic biology Cameron & Collins (2014), and opens the door to the exploration of novel dynamical behaviour when protein degradation is coupled to other synthetic circuits Cookson et al. (2011), Prindle et al. (2014). The most interesting bifurcation diagram found is the isola, which not only could be useful for error checking of deployed biosensors, but could be useful in clinical applications, for example *in vivo* detection of inflammation (Riglar et al. 2017) or metabolite levels (Rutter et al. 2019, 2021). Finally, the minimal mushroom topologies identified reveal a new way to create multistable systems, avoiding the need for loading a single bistable switch with additional autoregulation (Lou et al. 2010, Wu et al. 2017, Leon et al. 2016).

Our results complement the small amount of other existing work on the mushroom bifurcation. A recent study explored four incoherent feedforward networks with positive auto-regulation for their potential to display mushroom dynamics (Giri & Kar 2021). They found that all were capable of mushroom-like behaviour, though the appearance depended on the incoherence of the networks and the strength of positive feedback. Their systems contained the input signal on only one node, which is in contrast to our screening that suggested the double activating role of the input is a requirement to reproduce the mushroom diagram. This difference could be due to the assumptions on the underlying dynamics, since we assumed functions close to the Shea-Ackers formalism while Giri & Kar (2021) assumed an AND type logic gate interaction in the activation and repression of genes. This reinforces how the results of such computational investigations depend somewhat on the modelling choice of the dynamics, a result demonstrated directly previously Barnes et al. (2011). This suggests that further work in the experimental realisation of these systems is required to understand the actual dynamics, and how these depend on the particular biological context.

Overall, designing biological systems based directly on bifurcation properties provides a natural tool to explore phenotype-genotype relationships, relating topologies of the network with the available dynamical behaviours. Such an approach allows for the specification of a set of functional requirements without precising of explicit integration of the differential equations. In addition it provides a tool to design target dynamics that occur in a robust manner making these approaches key to the future of engineering of biological systems where uncertainty dominates.

## 4 STAR Methods

### 4.1 Gene Network Mixed-Integer modeling framework

The dynamics of gene regulation for the superstructure in Fig. 1 B is encoded using a mixed-integer framework based on deterministic ordinary differential equations. The gene circuit topology is characterized by a vector y of 9 integer variables (*y*_*uu*_, *y*_*vu*_, *y*_*wu*_ *y*_*uv*_, *y*_*vv*_, *y*_*wv*_, *y*_*wv*_, *y*_*ww*_) such that *y*_*ij*_ = −1 if *j* is repressed by *i, y*_*ji*_ = 1 if *i* is activated by *j*, and *y*_*ij*_ = 0 otherwise with *i, j* = {*u, v, w*}. The full network is characterized by the vector *y* and a vector *x* containing 12 real variables coding for tunable biochemical parameters (including promoter strengths, leakiness, degradation rate constants, repression and activation affinities). For example, the three gene system in Fig. 1 A (for which *y*_*uu*_ *= y*_*wu*_ *= y*_*vv*_ *= y*_*uw*_ *= y*_*ww*_ = 0 and *y*_*vu*_ *= y*_*uv*_ *= y*_*wv*_ *= y*_*wv*_ = −1) is given by:

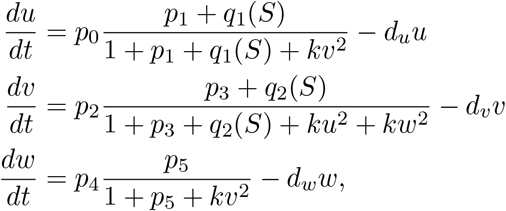

where the state vector is denoted by ω = (*u, v, w)* being *u, v, w* the concentrations of the proteins expressed by genes *U, V, W* respectively; *p*_0_, *p*_2_, *p*_4_ are the promoter strengths, *p*_1_*, p*_3_, *p*_5_ the promoter leakiness, k are repression strengths and *d*_*u*_, *d*_*v*_, *d*_*w*_ the degradation rate constants. The functions *q*_*1*_*(S), q*_*2*_*(S)* represent the concentration of activating transcription factors regulated by an input biochemical signal S, and are given by Hill functions

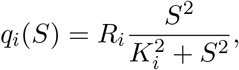

where *R* is the total concentration of transcription factor, *S* is the concentration of signal inducer, *K* is the dissociation constant, and the cooperativity is 2. We assume that repressor proteins bind as dimers (or equivalently there are two operator sites). The dynamics is expressed then in a compact form in terms of the vectors !, x, v and the input parameter S in this form:

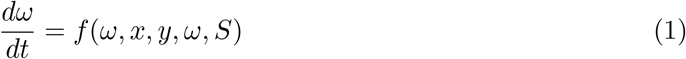

The system’s dimension (determined by the number of active genes) is in this case *N* = 3 (or *N* = 2 for the 2-gene case).

Depending on the specific design scenario, a number of assumptions can be made to reduce the parameter space dimensionality. In this work we assume *p*_3_ = *p*_1_ and *du* = 1.

### 4.2 conditions

To enable the optimization-based design of circuits with pre-specified bifurcation behavior (showing the target bifurcations at pre-specified locations in the extended parameter-state space), first we need to formulate the conditions for bifurcations in terms of the variables and parameters of the system: *φ, x, y, S*. In case of the mushroom-shaped diagram, the bifurcation of interest is the saddle-node bifurcation (also denoted as fold or limit point bifurcation). In particular, there are 4 saddle-node bifurcations (as indicated in Fig. 6 A). A parametric condition for saddle-node bifurcation in biochemical networks with mass action kinetics is demonstrated by Otero-Muras et al. (2012, 2017) and later generalized to arbitrary kinetics in Otero-Muras & Banga (2018). This condition, codified in the previously introduced mixed integer framework can be used for the direct design of systems with saddle-node bifurcations. Starting from Eq. 1 we compute the following extended Jacobian:

**Figure 6:**
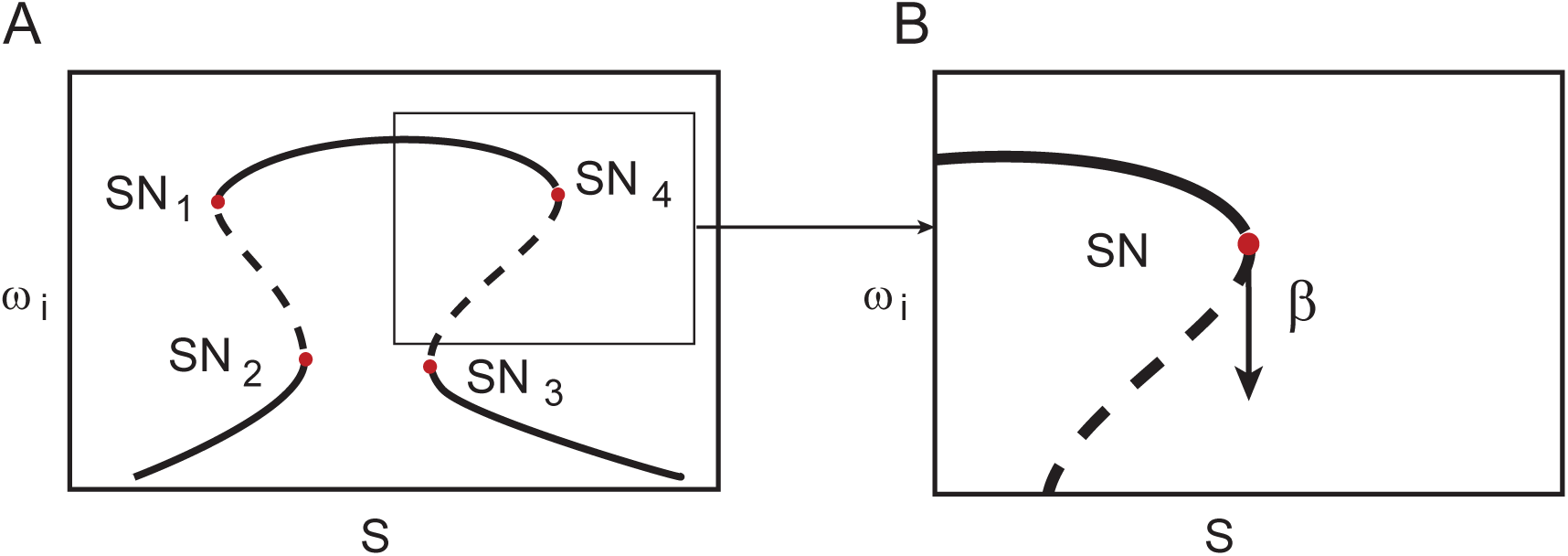
A) Mushroom-shaped bifurcation diagram and B) Saddle-Node bifurcation (the tangent vector (*β* at the bifurcation point is indicated).

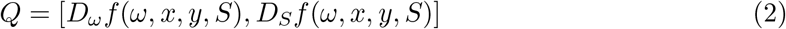

with dimensions *N × N* + 1. The tangent vector at the bifurcation point can be computed as (*β* = *null(Q)*, and the saddle-node occurs if (*β* _*N+1*_ = 0 (i.e., if the *N* + 1 entry of the tangent vector (*β* is zero). Therefore, we can define the function:

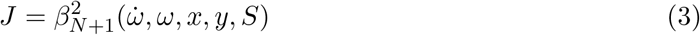

such that a saddle node is achieved when the function *J* reaches its minimum.

### 4.3 Optimization-based automated design

In order to design gene circuit with mushroom-shaped bifurcation diagrams we define the following objective function:

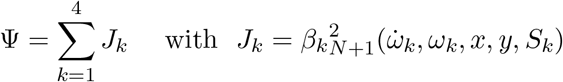

which takes value zero when four saddle-node bifurcations take place (in locations specified in the constraints). We then formulate the circuit design as the following optimization problem:

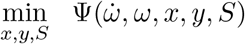

subject to:

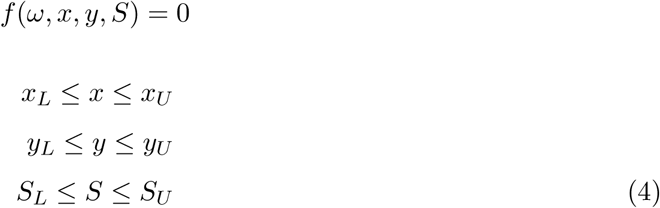

which is a mixed-integer nonlinear programming problem (MINLP). The nature of the problem is non-convex, therefore global optimization solvers are needed to obtain adequate solutions (Sendin et al. 2010). The solution of this optimization problem is not unique, since there are multiple combinations of topology and parameters leading to mushroom bifurcations. A single run of the algorithm will find a particular solution very efficiently (in the order of a few seconds for a standard PC), whereas a multistart strategy (running the optimization algorithm multiple times) will effectively find different topologies with mushroom bifurcation diagram. For the screening of 2-gene and 3-gene topologies with mushroom behaviour, we use a multistart strategy with 2000 and 10000 runs, respectively. As indicated in the Results section, the corresponding bounds for the parameters for the 2-gene and 3-gene networks are included in the supplemental tables S1 and S2, and the solution sets denoted by *Mushroom2D* and *Mushroom3D*, respectively.

The files with the codes and results (including *Mushroom2D* and *Mushroom3D*) are available online at DOI: 10.5281/zenodo.6024249.

In advanced design applications, it is usually the case that multiple objectives need to be taken into account (Otero-Muras & Banga 2017). For example, during the design of next generation biosensors, we might be interested not only in a mushroom-shaped diagram, but also in maximizing the regions of bistability. In numerous occasions, it is also required to operate at low protein burden to avoid compromising the viability of the cells, being the minimal protein production cost another important objective to be considered. In order to find the best designs with respect to multiple criteria (*Ψ*_1_,…, *Ψ*_*M*_), we consider a multi-objective formulation of the optimization problem., i.e. we set the mushroom bifurcation condition Ψ= 0 as a constraint of the following Multiobjective Mixed Integer Nonlinear Programming (MO-MINLP) problem:

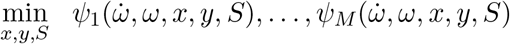

subject to:

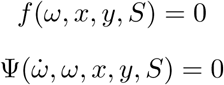

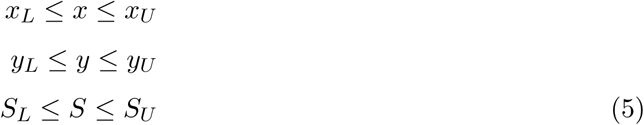

The solution of the above multicriteria mixed-integer nonlinear problem is not unique, but a set of vectors representing the best compromises between the (usually conflicting) objective functions. This set of best trade-offs is usually known as the Pareto set.

#### 4.3.1 ptimization strategy and solvers

Computing the Pareto set of optimal designs by solving the above problems efficiently and reliably can be a daunting task due to their non-convexity, arising from their highly constrained, partially discrete and non-linear nature.

Here we transform the multicriteria mixed-integer formulation (MO-MINLP) into a finite set of single-objective mixed-integer (MINLP) problems by adopting an “-constraint approach (Miettinen 2012). The resulting set of MINLPs is then solved using a hybrid strategy, eSS-MISQP, which combines a diversification phase (using a global optimization metaheuristic, eSS) with intensification steps (using an efficient local mixed-integer optimization solver, MISQP). This hybrid strategy has been found to outperform other global approaches in terms of both efficiency and robustness (Otero-Muras & Banga 2017).

#### 4.3.2 Analysis of Robustness

In order to assess the robustness of a topology, we use the metric defined in (Otero-Muras & Banga 2019) in terms of the interquantile ranges (IQR) measuring the spread of the distributions of each of the parameters for the solution set. To this aim, we build the corresponding robustness proxy histograms (see the supplemental Figure S6). The solution set *Mushroom3D* is obtained by solving the optimization problem (4) with a multistart strategy, and the parameters for the search are included in the supplemental Table 2. The robustness proxy histograms for the extremes of the Pareto Front Figure in Figure 4A are included in the supplemental Figure S6.

## 5 Acknowledgements

IOM and JRB acknowledge funding from MCIN/AEI/ 10.13039/501100011033 and “ERDF A way of making Europe” through grant DPI2017-82896-C2-2-R (SYNBIOCONTROL). JRB also acknowledges funding from MCIN/AEI/ 10.13039/501100011033 through grant PID2020-117271RB-C22 (BIODYNAMICS). RPC acknowledges financial support by the UCL Mathematics Clifford Fellowship. CPB received funding from the European Research Council (ERC) under the European Unions Horizon 2020 research and innovation programme (Grant No. 770835) and from the Wellcome Trust (209409/Z/17/Z).

## Supplemental Information

**Table S1:** Upper and Lower bounds (UB and LB respectively) for the decision variables of the optimization problem, including the parameters and the states at the bifurcation points S1 and S2 (2 dimensional gene regulatory network)

| | $p_0$ | $p_1$ | $p_2$ | $k$ | $R_1$ | $R_2$ | $dv$ | $K1$ | $K2$ | $S_1$ | $S_2$ |
| --- | --- | --- | --- | --- | --- | --- | --- | --- | --- | --- | --- |
| UB | 300 | 300 | 300 | 0.1 | 300 | 300 | 10 | 150 | 150 | 50 | 200 |
| LB | 50 | 50 | 50 | 0.01 | 10 | 10 | 0.1 | 10 | 10 | 5 | 80 |
| | $u(S_1)$ | | $v(S_1)$ | | $u(S_2)$ | | $v(S_2)$ | | | | |
| UB | 100 |  | 200 |  | 100 |  | 200 |  |  |  |  |
| LB | 50 |  | 10 |  | 50 |  | 10 |  |  |  |  |

**Table S2:**
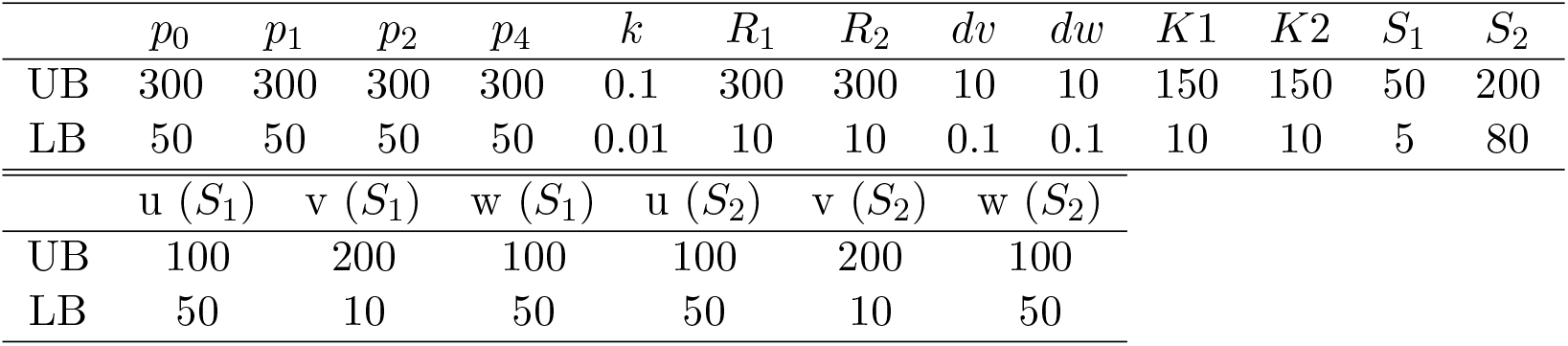
Upper and Lower bounds (UB and LB respectively) for the decision variables of the optimization problem, including the parameters and the states at the bifurcation points S1 and S2 (3 dimensional gene regulatory network)

**Table S3:**
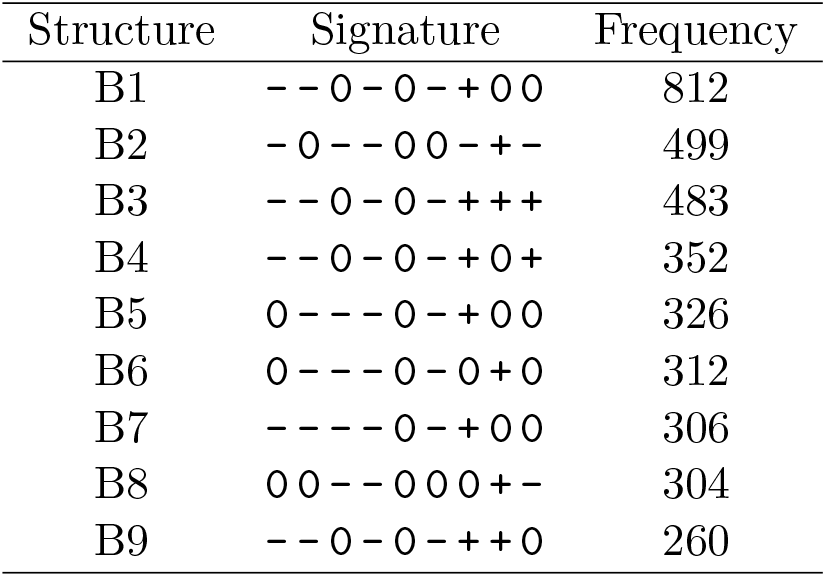
Most frequent 3-gene topologies obtained with the robustness analysis. The corresponding graphs can be found in Fig. S1

**Table S4:**
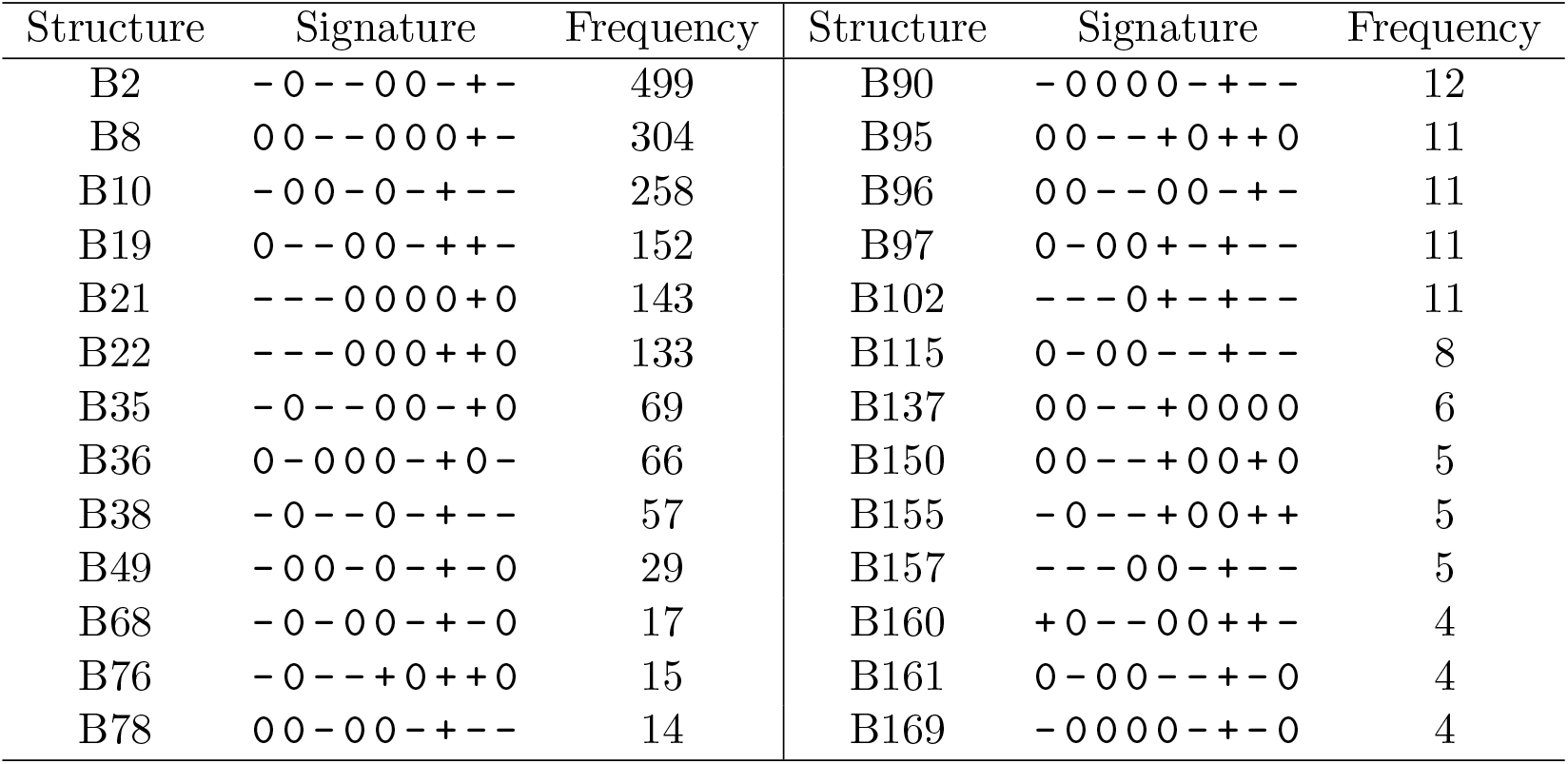
Topologies of 3-gene regulatory networks showing mushroom bifurcation behaviour (and that, additionally, are not built up from 2-gene mushroom structures). Graphs for the first 9 topologies are depicted in Fig. S2

**Table S5:**
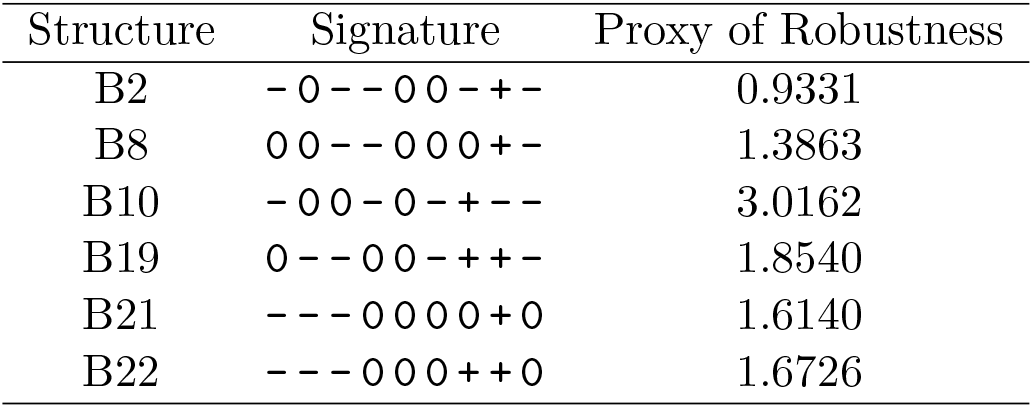
Proxy of Robustness for the first 5 structures in Table S4 (the corresponding topologies are depicted in Fig. S2)

| Structure | Signature | Proxy of Robustness |
| --- | --- | --- |
| B2 | -0--00-+- | 0.9331 |
| B8 | 00--000+- | 1.3863 |
| B10 | -00-0-+-- | 3.0162 |
| B19 | 0--00-++- | 1.8540 |
| B21 | ---0000+0 | 1.6140 |
| B22 | ---000++0 | 1.6726 |

**Figure S1:**
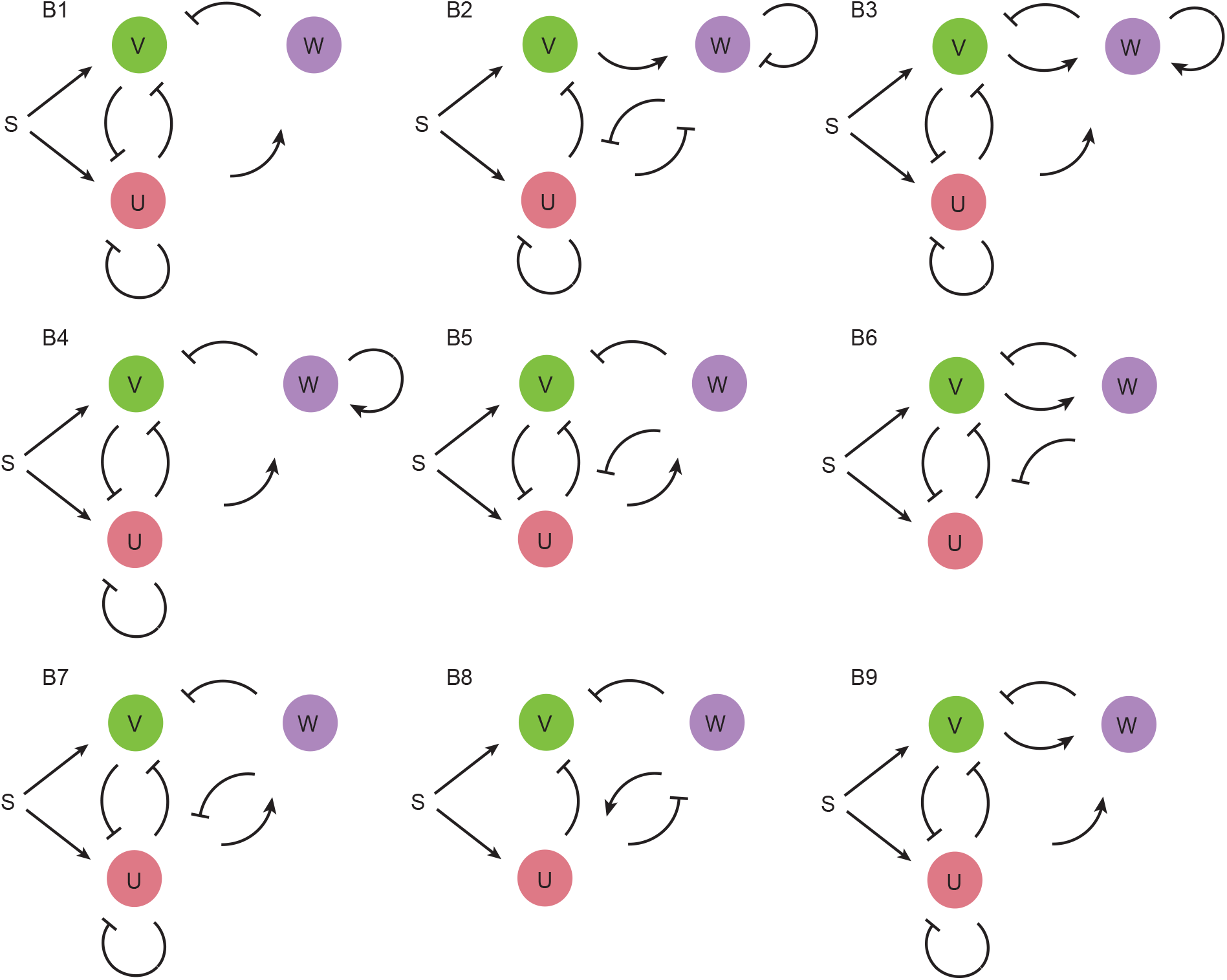
Most frequent 3 gene topologies (Table S3)

**Figure S2:**
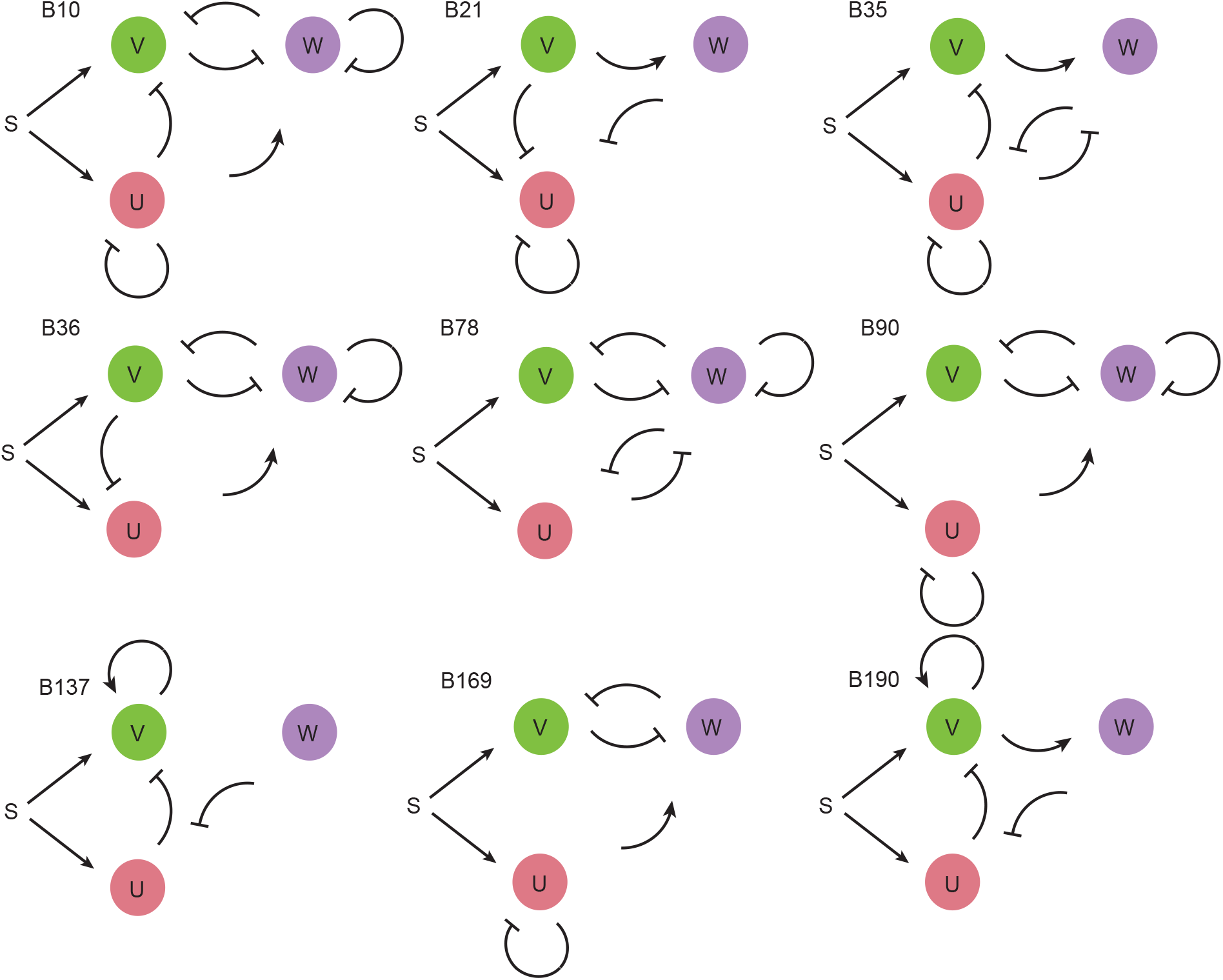
Selection of 3-gene topologies (Table S4)

**Figure S3:**
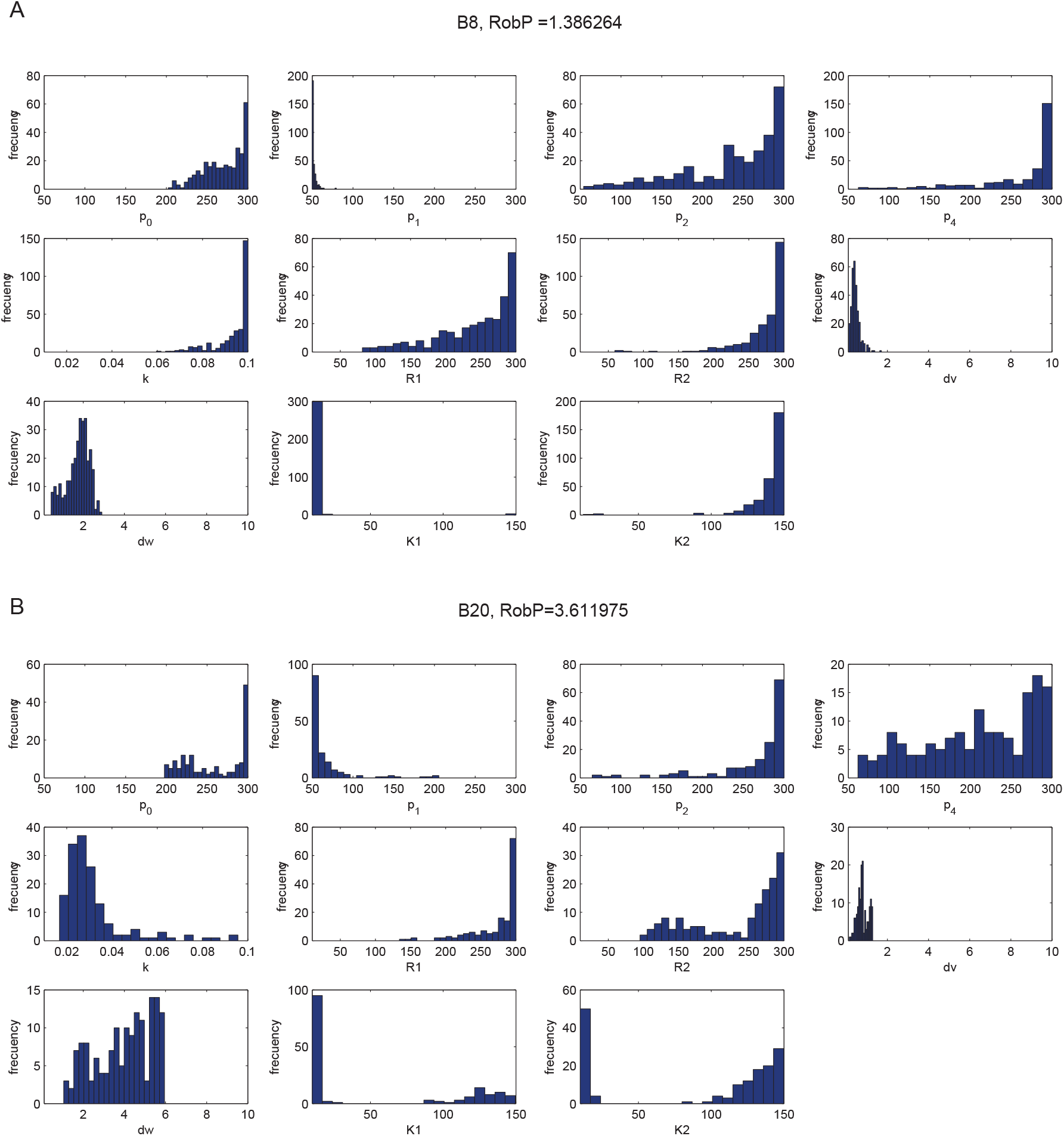
Robustness proxy histograms for the two extremes of the Pareto Front in the main text Fig. 3 B. The minimum and maximum x-axis values are the lower and upper bounds for the parameters, respectively

